# Harnessing Smartphone RGB Imagery and LiDAR Point Cloud for Enhanced Leaf Nitrogen and Shoot Biomass Assessment - Chinese Spinach as a Case Study

**DOI:** 10.1101/2024.10.26.620395

**Authors:** Aravind Harikumar, Itamar Shenhar, Miguel R. Pebes-Trujillo, Lin Qin, Menachem Moshelion, Jie He, Kee Woei Ng, Matan Gavish, Ittai Herrmann

## Abstract

Accurate estimation of leaf nitrogen concentration and shoot dry-weight biomass in leafy vegetables is crucial for crop yield management, stress assessment, and nutrient optimization in precision agriculture. However, obtaining this information often requires access to reliable plant physiological and biophysical data, which typically involves sophisticated equipment, such as high-resolution in-situ sensors and cameras. In contrast, smartphone-based sensing provides a cost-effective, manual alternative for gathering accurate plant data.

In this study, we propose an innovative approach for estimating leaf nitrogen concentration and shoot biomass by integrating smartphone RGB imagery with Light Detection and Ranging (LiDAR) data, using *Amaranthus dubius* (Chinese spinach) as a case study. The influence of varying nitrogen dosages on individual spectral and structural features derived from smartphone RGB imagery and LiDAR data was modeled. Additionally, the spectral indices from RGB imagery and structural indices from LiDAR data were combined to model both leaf nitrogen concentration and shoot biomass.

The performance of crop parameter modeling was evaluated using support vector regression, random forest regression, and lasso regression. Results demonstrate that the combined use of smartphone RGB imagery and LiDAR data can accurately estimate leaf total reduced nitrogen concentration, leaf nitrate concentration, and shoot dry-weight biomass, with average relative root mean square errors as low as 0.06, 0.16, and 0.05, respectively. Furthermore, the optimal nitrogen dosage for maximizing biomass yield in Chinese spinach was also estimated using the smartphone data. This study lays the groundwork for smartphone-based estimate leaf nitrogen concentration and shoot biomass, supporting accessible precision agriculture practices.

## 1 Introduction

Nitrogen is a cornerstone mineral nutrient essential for regulating the yield and overall health of crop plants, as it is a key component of plant amino acids, proteins, and nucleic acids. Subtropical leafy vegetable crops, such as *Amaranthus dubius* (Chinese spinach), are experiencing a growing global market demand due to their exceptional nutritional profile and relatively affordable price [1]. These crops respond notably well to nitrogen fertilization, attributed to their efficient nitrogen uptake mechanisms paired with less efficient nitrogen reductive systems [2, 3]. However, the nitrogen fertilizer use efficiency for leafy vegetable crops typically remains below 50% of the applied amount [4, 5], reducing profitability and contributing to substantial nitrogen loss to the environment. Such loss leads to increased nitrogen runoff into water bodies and elevated greenhouse gas emissions [6]. Balancing crop nitrogen needs with fertilizer application is thus critical to optimize crop performance, enhancing nutritional quality, protecting the environment, and maximizing returns for farmers. Given nitrogen’s vital role in improving both nutritional value and market appeal of leafy vegetables, accurate estimation of leaf nitrogen concentration and shoot dry-weight biomass is essential for determining the specific nitrogen requirements of crop species and, consequently, for optimizing fertilizer application to maximize yield and profitability.

Traditional methods for assessing leaf nitrogen concentration, such as laboratory-based micro Kjeldahl analysis [7], and for determining shoot dry-weight biomass through drying, rely on destructive sampling, making them impractical and costly in terms of time, financial resources, and labor. This necessitates the development of innovative approaches that allow timely, non-destructive estimation of these plant parameters. Smartphone sensing provides a non-destructive method for analyzing crop dynamics, representing a promising pathway to high-throughput plant phenotyping that can transform crop management practices [8]. Extensive research has consistently demonstrated a robust relationship between Red-Green-Blue (RGB) imagery and leaf nutrient composition [9, 10]. For example, studies indicate that leaf chlorophyll concentration, which correlates positively with leaf nitrogen levels, provides a reliable indicator of nitrogen concentration [11]. The emergence of optical smartphone RGB imaging has thus provided a cost-effective approach for capturing crop traits compared to the use of expensive cameras. The widespread availability and affordability of smartphones have driven the development of methods that use smartphone-based RGB imagery to estimate leaf nitrogen concentration in staple crops, such as rice and wheat [12, 13]. However, a key limitation of RGB imagery is its lack of structural data, which is needed to estimate traits like crop plant height and shoot dry-weight biomass [14].The recent integration of Light Detection and Ranging (LiDAR) sensors into smartphones marks a groundbreaking advancement, enabling the collection of structural plant data, including biomass estimation. Combining smartphone RGB data with LiDAR data offers potential for estimating both biophysical and physiological traits in leafy vegetable crops. For instance, Bar-sella et al. successfully estimated leaf transpiration in maize using smartphone RGB imagery and LiDAR data collected with an iPhone 13 Pro Max smartphone [15].

A comprehensive literature review reveals a significant research gap in the integration of relatively low-cost smartphone RGB and LiDAR data for estimating plant traits, including leaf nitrogen concentration and shoot biomass. Moreover, few studies have focused on nitrogen concentration and biomass estimation specifically for high-demand tropical leafy vegetables, such as Chinese spinach. Additionally, many of the existing studies have been conducted in open-field conditions, which are subject to considerable variations by biotic and abiotic factors [15, 16], whereas the current study focuses on semi-controlled tropical greenhouse conditions. Specifically, this study aims to achieve three main objectives: (a) to evaluate the effectiveness of combined smartphone-based RGB imagery and LiDAR data for accurately estimating leaf nitrogen concentration and shoot biomass in leafy vegetable crops within a semi-controlled greenhouse setting, (b) to quantify the relevance of various RGB and LiDAR-based vegetation indices for estimating leaf nitrogen concentration and shoot dry-weight biomass in Chinese spinach, and (c) to identify the optimal nitrogen dosage for maximizing crop dry-weight biomass yield using integrated smartphone RGB imagery and LiDAR data.

## 2 Materials

### 2.1 Study Site, Experimental Setup and Crop Management

The experiment was conducted in a semi-controlled tropical greenhouse facility that is (Oasis Living Lab, Singapore), specifically dedicated to leafy vegetable production, from September to November 2023. A total of 84 Chinese spinach plant species, known for their high productivity and nutritious value, were selected for this study (Fig. 1). Seeds were initially sown in seedling trays and nurtured for 3–4 weeks to ensure optimal growth prior to transplantation [5]. At the appropriate growth stage, the plants were transplanted into 4-L pots (Tefen Ltd., Nahsholim, Israel) filled with coco-peat substrate (Riococo Ltd., Sri Lanka) at the experiment’s commencement.

**Figure 1:**
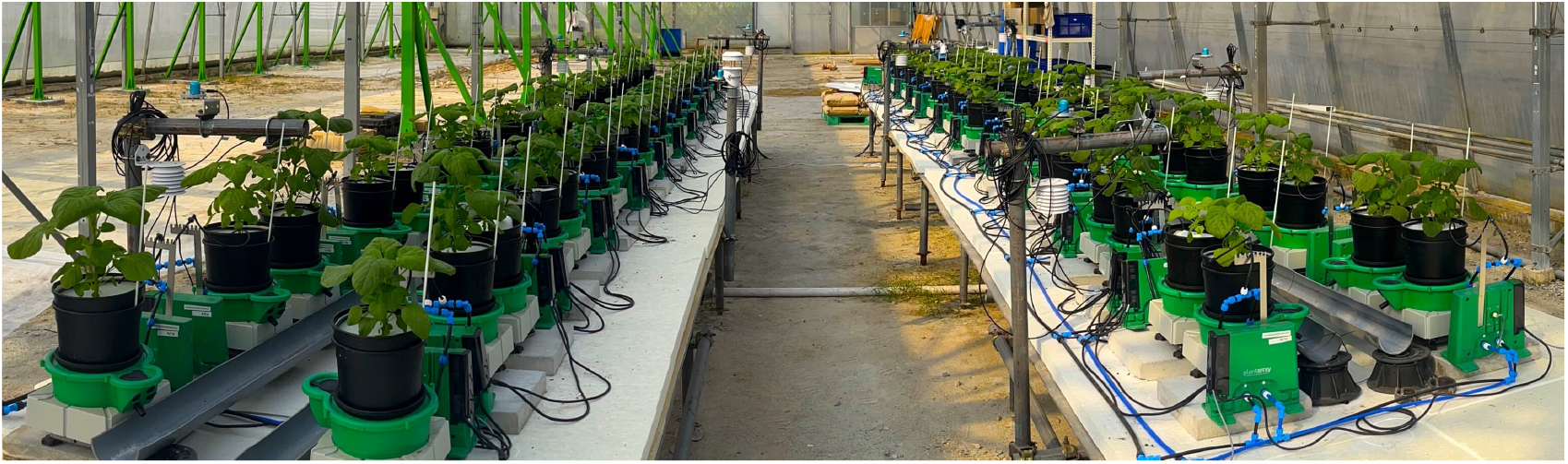
Experimental setup of 84 Chinese spinach plants in the semi-controlled greenhouse at the OasisLab, Singapore. Nitrogen dosing for each plant is precisely regulated using the PlantArray system, allowing controlled assessment of varying nitrogen treatments (in randomized block design) on plant growth and performance.

To assess the impact of varying nitrogen dosages on plant growth and performance, plants were divided into eight distinct treatment groups, each with 10 or 11 replicates. Each group received one of the following nitrogen dosages: 20, 40, 80, 120, 160, 200, 400, or 800 parts per million (ppm), administered in an aqueous solution. Nitrogen application was meticulously managed using precision pumps (Shurflo 5050-1311-H011, Pentair, MN, USA) under the control of the PlantArray system (Plant-Ditech, Yavne, Israel), as described in [17].

At the conclusion of the experiment, plant shoots were harvested and manually separated into leaves and stems. The harvested shoots were then dried in an oven at a constant temperature of 60°C for one week to ensure thorough desiccation before laboratory analysis. These dried samples were analyzed to determine leaf total reduced nitrogen (*TRN*) concentration, leaf nitrate (*NO*_3_) concentration, and shoot dry-weight (*DW*) biomass. Here, *TRN* represents the total nitrogen content in its reduced forms, including nitrogen incorporated into amino acids, proteins, and other organic compounds, while *NO*_3_ denotes the inorganic nitrogen absorbed by plants from the soil.

### 2.2 Smartphone Data Collection

In this study, an iPhone 14 Pro Max smartphone (Apple, Cupertino, CA, USA) with a field of view (FoV) of 69°, pixel size (PL) of 1.9 *µ*m, and focal length (F) of 26 mm was mounted on a manually held ZHIYUN Smooth 5 gimbal platform (Guilin Zhishen Information Technology Co., Ltd., Shenzhen, China) to acquire RGB imagery from a nadir perspective (Fig. 2a). RGB images for all 84 Chinese spinach plants were captured within a 10-minute timeframe under clear skies, between 11:30 am and 12:00 pm, on the day preceding harvest. The RGB data acquisition height was set to ensure the entire plant crown fell within the camera’s field of view, specifically at 50 *±* 5 cm above the highest point of the plant crown. Precise horizontal alignment and smartphone orientation during data capture were verified using the Pocket Bubble Level iPhone application (ExaMobile S.A., Bielsko-Biala, Poland) to maintain a level device position throughout the imaging process. At the end of data collection, an RGB image of a 99.9% circular white reference panel was taken to facilitate the conversion of raw RGB digital numbers to reflectance values.

**Figure 2:**
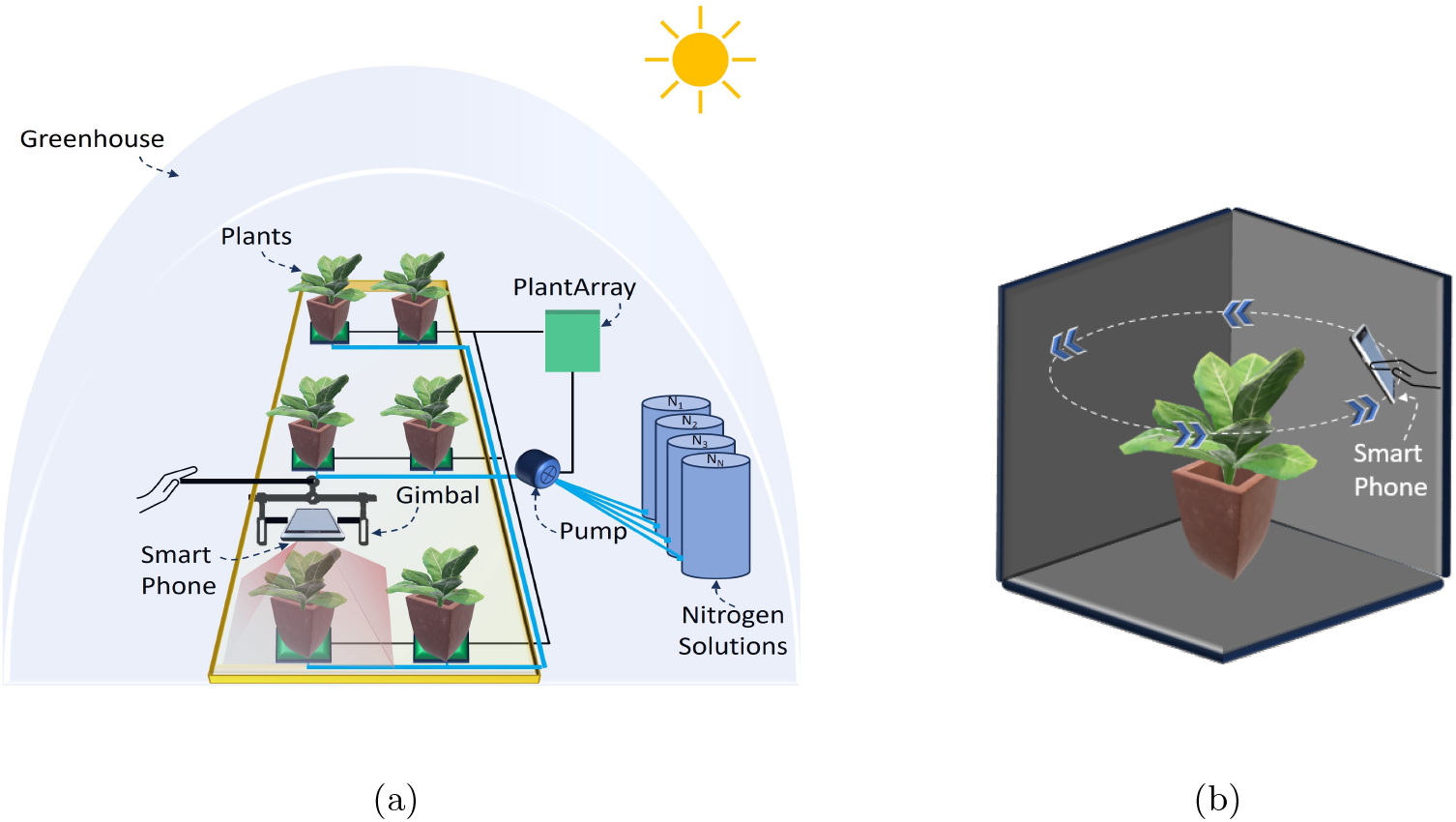
Schematic representation of the smartphone-based data acquisition setup in the indoor greenhouse. (a) RGB imagery acquisition setup showing the smartphone mounted on a gimbal platform and the PlantArray system managing nitrogen dosing for individual plants. (b) 3D LiDAR point cloud data acquisition setup, where the smartphone is maneuvered around the plant to capture detailed structural data of the plant crown

Additionally, 3D LiDAR point cloud data of each plant was captured using the iPhone 14 Pro Max’s in-built LiDAR sensor, operated via the Polycam 3D scanning application (Polycam, Inc., San Francisco, California.). The LiDAR sensor specifications are consistent across iPhone 13 and iPhone 14 Pro models, allowing them to be used interchangeably for 3D data acquisition. Plants were positioned against a dark background, 10–20 cm from the pot, to reduce background noise and enhance point cloud quality. The Polycam application’s LiDAR capture mode collected high-density point cloud data, achieving a resolution exceeding 100 points per square centimeter. During LiDAR data acquisition, the smartphone was maneuvered in a circular path around and above each plant (Fig. 2b) to obtain a comprehensive, detailed representation of the plant’s crown structure. The Polycam application generated the final raw 3D point cloud, with each point containing R, G, and B value attributes, within 20–30 seconds of scan completion and preprocessing initiation.

## 3 Methods

The proposed approach characterizes physiological and biophysical crop traits by leveraging spectral and structural data obtained from smartphone RGB imagery and LiDAR point cloud data (Fig. 3). RGB imagery processing includes generating reflectance values to account for environmental variations in solar illumination, followed by leaf segmentation and extraction of spectral-reflectance features. Structural feature extraction from LiDAR data involves denoising and segmenting the 3D point cloud to isolate plant data from background objects and noise.

**Figure 3:**
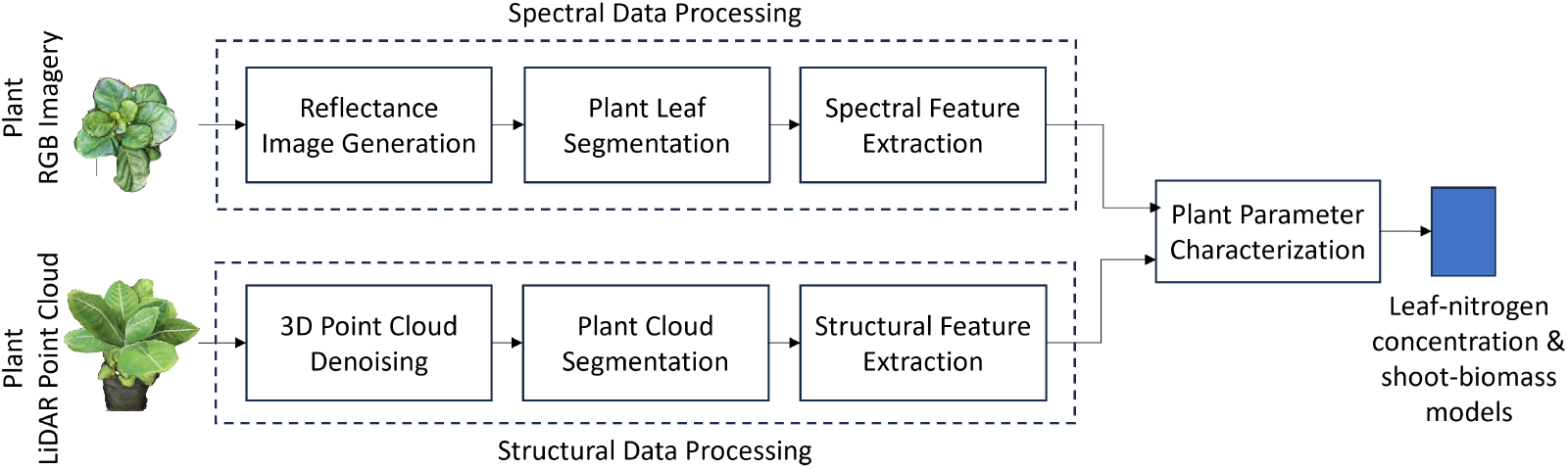
Block diagram of the smartphone-based leaf nitrogen concentration and shoot dry-weight biomass estimation setup for Chinese spinach. Spectral features derived from smartphone RGB imagery and structural features from LiDAR point cloud data are combined to estimate key leaf nitrogen concentration and shoot biomass.

## 3.1 Spectral Data Processing

The inherent sensor-electronic noise present in the measured values was mitigated by performing dark image subtraction on the raw optical smartphone RGB images. A dark image was generated by covering the smartphone’s rear camera with a dark cloth. The resulting RGB images were then converted to reflectance values by dividing each digital number (DN) value by the average DN value from a 99.9% white reference image [18].

The individual RGB reflectance images captured during our experiment contained extraneous details, including background objects such as pots and other experimental hardware (Fig. 4a). To minimize estimation errors, accurate delineation of plant leaves from the background was essential. We employed the Mask Convolutional Neural Networks (MaskCNN) algorithm [19] for automatic leaf segmentation, chosen for its precision in identifying leaf structures (i.e., the objects of interest) within images. The CNN was trained on 245 individual leaf images, each carefully annotated by an expert from 150 randomly selected plant images. To enhance dataset size, diversity, and robustness, data augmentation techniques—such as rotations (45^°^, 90^°^ and 135^°^) and scalings (0.10, 0.25, 0.50, 0.75, and 1.25) - were applied to each original image, resulting in a total of 2,250 leaf samples for model training. The MaskCNN algorithm used distinct leaf traits such as shape, texture, and color to automatically extract leaf boundaries from the background (Fig. 4b). The output of the MaskCNN segmentation process was a mask layer overlaying the original RGB reflectance image, where pixels corresponding to plant leaves were assigned a value of 1 and background pixels a value of 0. This binary mask effectively isolated the leaf regions from the background by multiplying it with the vegetation spectral index image, providing a clean extraction of the plant leaves in the vegetation spectral index imagery (Fig. 4c).

**Figure 4:**
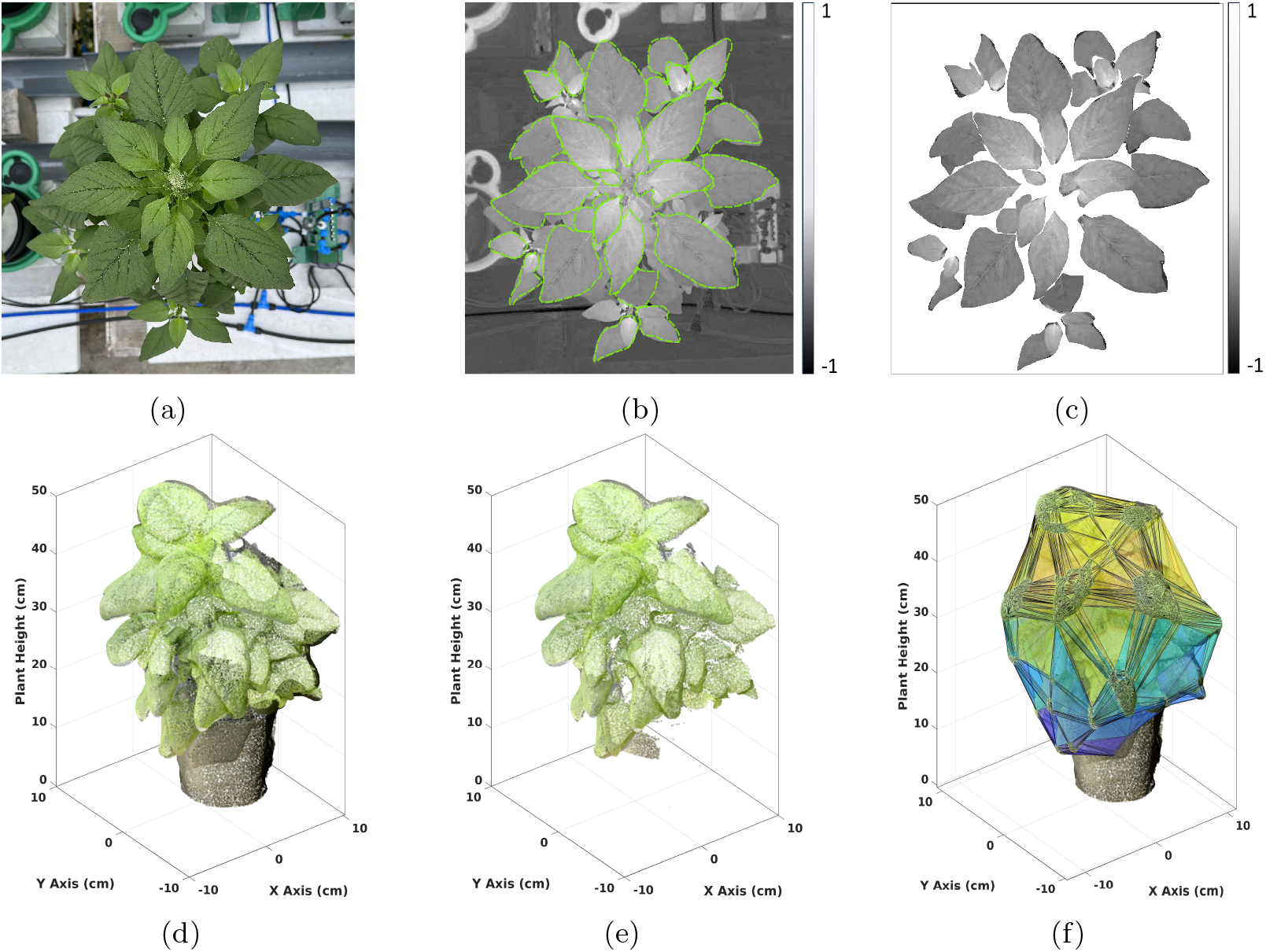
Visualization of Data Processing: (Top row:) Top view (a) RGB image of a Chinese spinach plant captured using a smartphone. (b) Spectral index image, with boundaries delineated using Mask-CNN for leaf segmentation. (c) Resulting leaf-segmented image. (Bottom row:) (d) 3D LiDAR point cloud of the plant, (e) filtered 3D point cloud after K-means clustering to isolate plant data, and (f) convex hull fitted to the filtered plant point cloud to extract structural features.

To assess the accuracy of the MaskCNN-based leaf delineation, we utilized the Intersection over Union (IoU) metric (see Equation (1)). This metric compares the estimated leaf mask against a manually delineated leaf mask. Let *A*^*est*^ denotes the estimated leaf-mask area and *A*^*ref*^ denotes the manually delineated leaf-mask area, then IoU is estimated as

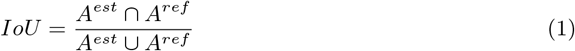

An IoU value of 1 indicates perfect overlap, while a value of 0 indicates no overlap [20]. MaskCNN segmentation performance was assessed using a set of 245 manually extracted leaves from 25 randomly selected plants across all nitrogen fertigation treatments.

The leaf regions identified by the MaskCNN segmentation algorithm were then extracted from the RGB reflectance images to estimate vegetation indices. These segmented leaf regions were used to calculate various vegetation indices that serve as proxies for different plant traits. The RGB spectral indices were quantitative measures derived from plant leaf reflectance values, reflecting their sensitivity to specific plant characteristics, such as chlorophyll content and overall vegetation vigor.

Chlorophyll concentration in leaves is a critical parameter for assessing plant greenness, as it reflects photosynthetic capacity, growth vigor, and shoot biomass [21, 22]. The Excess Green Index (ExG), Green Leaf Index (GLI), and Normalized Green-Red Difference Index (NGRDI) are commonly used to evaluate chlorophyll content in leaves [23, 24]. The ExG index, calculated by subtracting the red and blue reflectance from twice the green reflectance, is highly responsive to changes in chlorophyll concentration. The GLI measures vegetation “greenness” by assessing green reflectance relative to other wavelengths [25]. Similarly, the NGRDI, which normalizes the difference between green and red reflectance, also serves as an indicator of chlorophyll concentration.

In addition to chlorophyll content, certain indices are designed to evaluate other leaf characteristics, such as pigmentation, stress, and structure. For example, the Triangular Greenness Index (TGI) [26] and Normalized Difference Yellowness Index (NDYI) [27] are valuable in this regard. Healthier leaves often have certain thicknesses and internal structures that influence light absorption and reflection; therefore, variations in TGI values can indicate differences in leaf morphology, such as thickness or density, affecting light interaction [26]. The NDYI, which measures the difference between green and blue reflectance, provides insights into yellow pigments, suggesting potential stress or senescence [27]. Both indices are instrumental for monitoring vegetation health and detecting stress conditions [28, 29].

In this study, we used a total of 15 widely recognized RGB spectral indices from the literature. Table 1 provides a detailed summary of these RGB spectral indices, including the parameters and calculations used to derive them from the smartphone-based RGB imagery. The average value of all leaf pixels were computed to obtain a plant-level RGB feature estimate for each index.

**Table 1:**
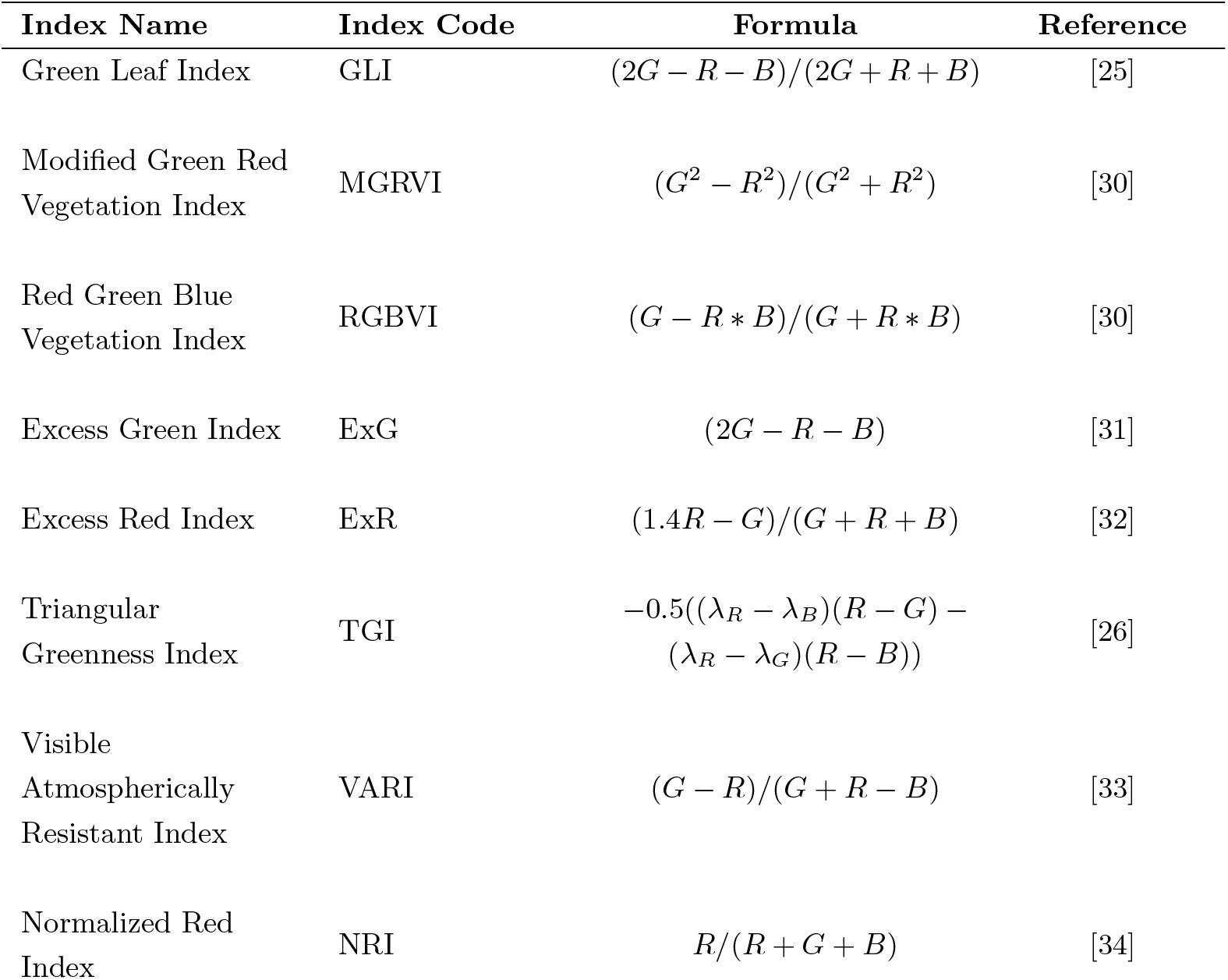
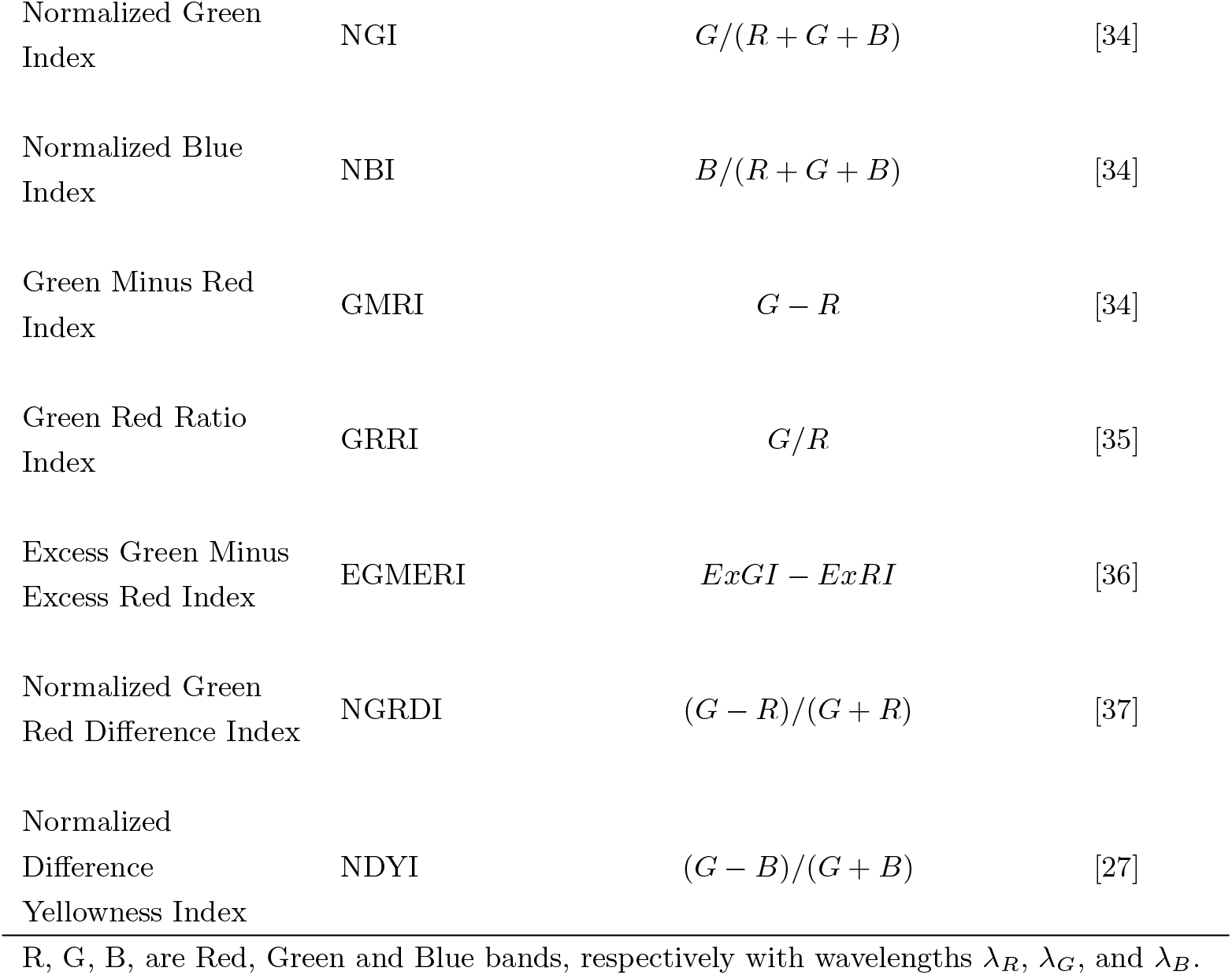
The spectral indices derived from the smartphone RGB imagery of plants.

| Index Name | Index Code | Formula | Reference |
| --- | --- | --- | --- |
| Green Leaf Index | GLI | $(2G - R - B)/(2G + R + B)$ | [25] |
| Modified Green Red Vegetation Index | MGRVI | $(G^2 - R^2)/(G^2 + R^2)$ | [30] |
| Red Green Blue Vegetation Index | RGBVI | $(G - R * B)/(G + R * B)$ | [30] |
| Excess Green Index | ExG | $(2G - R - B)$ | [31] |
| Excess Red Index | ExR | $(1.4R - G)/(G + R + B)$ | [32] |
| Triangular Greenness Index | TGI | $-0.5((\lambda_R - \lambda_B)(R - G) - (\lambda_R - \lambda_G)(R - B))$ | [26] |
| Visible Atmospherically Resistant Index | VARI | $(G - R)/(G + R - B)$ | [33] |
| Normalized Red Index | NRI | $R/(R + G + B)$ | [34] |
| Normalized Green Index | NGI | $G/(R + G + B)$ | [34] |
| Normalized Blue Index | NBI | $B/(R + G + B)$ | [34] |
| Green Minus Red Index | GMRI | $G - R$ | [34] |
| Green Red Ratio Index | GRRI | $G/R$ | [35] |
| Excess Green Minus Excess Red Index | EGMERI | $ExGI - ExRI$ | [36] |
| Normalized Green Red Difference Index | NGRDI | $(G - R)/(G + R)$ | [37] |
| Normalized Difference Yellowness Index | NDYI | $(G - B)/(G + B)$ | [27] |

R, G, B, are Red, Green and Blue bands, respectively with wavelengths $\lambda_R$ , $\lambda_G$ , and $\lambda_B$ .

### 3.2 Structural Data Processing

Let *P* = *{ρ*_*i*_ *∈* ℝ^3^ | *i* = 1, 2, …, *M}*, with *ρ*_*i*_ = (*x*_*i*_, *y*_*i*_, *z*_*i*_), be the 3D LiDAR point cloud obtained using the smartphone (Fig. 4d). Each point in the raw point cloud is composed of the three-dimensional space characterized by the x, y, and z Euclidean coordinates derived from the iPhone LiDAR sensor, together with the R, G, and B attribute values derived internally using the iPhone camera. However, the raw 3D point cloud often contains noisy points which arise from various sources such as suspended particles in the air, glitches in electronics, or errors in data acquisition and processing. To ensure accurate downstream analysis, it is essential to remove these noisy points from the point cloud. This denoising process involved several steps: a) voxelization: The volume covered by the 3D point cloud was divided into *S* regular cubic voxels *{V*_*i*_ | *i* = 1, 2, …, *S}*, each with sides of equal length (i.e., 1cm), b) point density (PD) calculation: The number of points contained within each voxel was calculated, c) points with a voxel PD below a certain threshold were identified as noisy points and subsequently removed from the point cloud. The threshold value was chosen carefully to minimize both omission (failure to detect noisy points) and commission (incorrectly identifying valid points as noisy) errors. In our case this threshold has been identified to be equal to 4 data points. By implementing this denoising procedure, the resulting point cloud was cleansed of isolated points, facilitating accurate structural parameter estimation (Fig. 4e).

The subset of points *G* = *{p*_*j*_ | *i* = 1, 2, …, *N}* ⊂ *P* which is part of the plant object was identified by using the two-class (i.e., *K* = 2) K-means clustering approach. This clustering method delineates the background data points from the plant data jointly using the R, G, and B attributes. By clustering the points into two distinct classes, one representing the black background and the other representing the green plant, we can effectively separate the vegetation from other objects in the scene (Fig. 4f). This segmentation process played a crucial role in isolating the plant canopy from the surrounding environment, enabling more accurate analysis and interpretation of the structural parameters derived from the LiDAR point cloud.

Structural attributes encapsulating the physiological characteristics of the plant were derived from *G*. These attributes encompass essential matrices such as plant height, maximum crown span, point density, and crown volume, providing valuable insights into the plant’s structural composition. The plant height (HT) was determined as the elevation of the highest point within the plant point cloud *G*. Meanwhile, the maximum crown span (CW), was calculated as the greatest distance between any two points within the point cloud *G* when projected onto the ground plane (X-Y plane), effectively capturing the lateral extent of the plant canopy. To assess the density of the crown (CD), we calculated the average point count within each 1cm voxel containing at least one point. This voxel-based approach provides a robust measure of crown density, offering insights into the spatial distribution of foliage within the plant canopy. By encapsulating the plant canopy within a convex envelope, a holistic estimate of the plant’s crown volume (CV) is obtained, facilitating a comprehensive understanding of the plant structure. The CV was estimated as the volume enclosed by the convex hull *V* ^*hull*^ formed around *G* (Fig. 4f). Here, the boundary fraction parameter *α* of the convex hull was set to 0.5 to realize a 3D hull which is neither too tight not too loose. The other structural traits included point count, point density and point density variance. The number of points above 75%, 50% and 25% of the plant height were represented as *N*_75_, *N*_50_ and *N*_25_, respectively. The 9 plant-level LiDAR structural features used in the study are defined in Table 2.

**Table 2:**
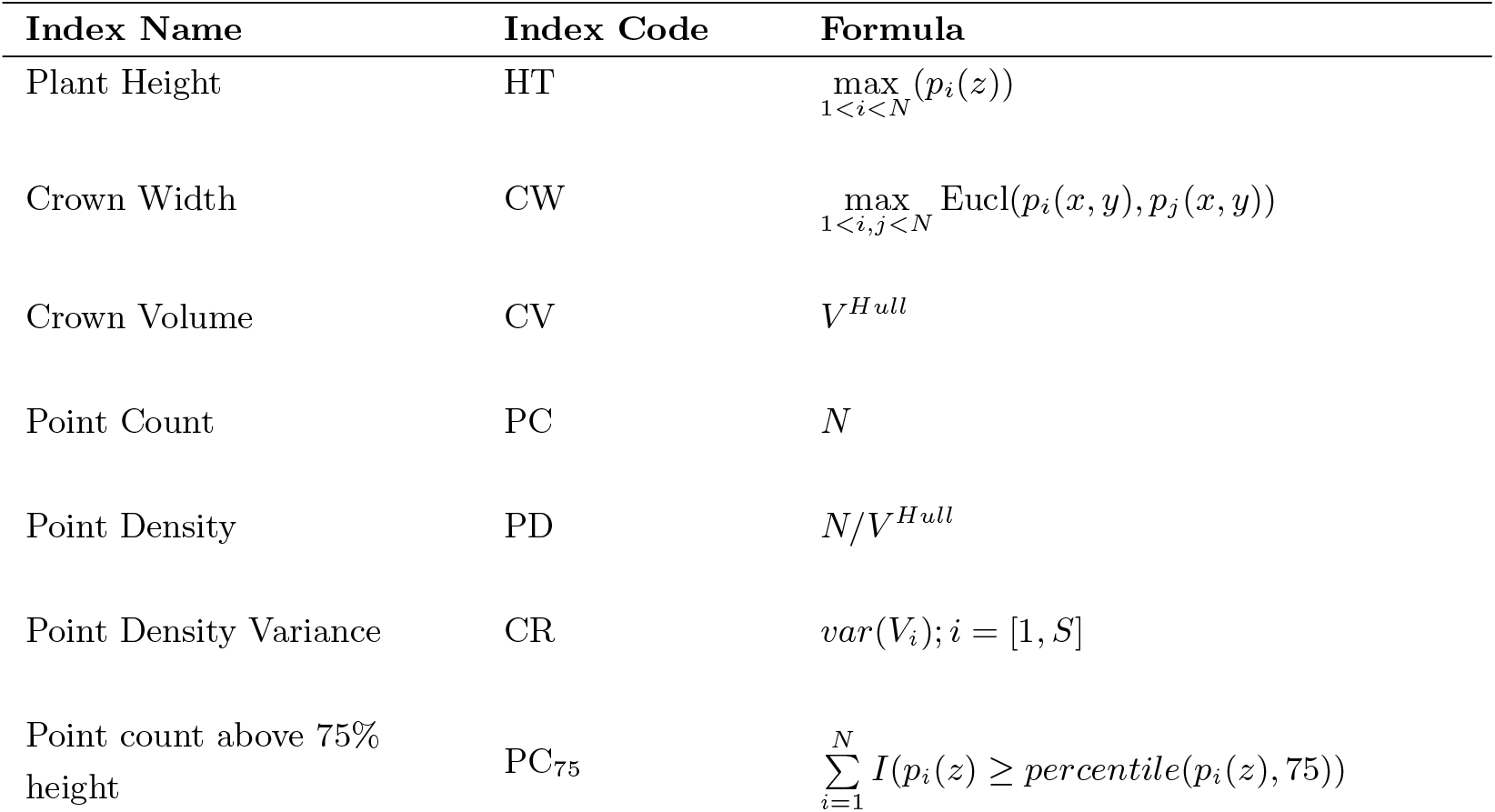
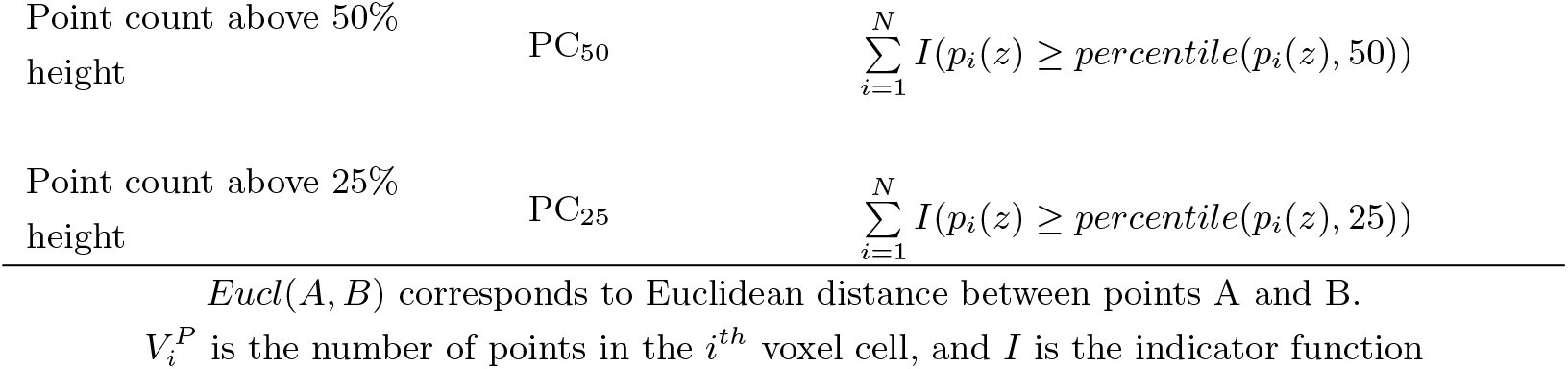
The structural indices derived from the smartphone LiDAR point cloud of plants.

| Index Name | Index Code | Formula |
| --- | --- | --- |
| Plant Height | HT | $\max_{1 \leq i \leq N} (p_i(z))$ |
| Crown Width | CW | $\max_{1 \leq i, j \leq N} \text{Eucl}(p_i(x, y), p_j(x, y))$ |
| Crown Volume | CV | $V^{Hull}$ |
| Point Count | PC | $N$ |
| Point Density | PD | $N/V^{Hull}$ |
| Point Density Variance | CR | $var(V_i); i = [1, S]$ |
| Point count above 75% height | $PC_{75}$ | $\sum_{i=1}^N I(p_i(z) \geq \text{percentile}(p_i(z), 75))$ |
| Point count above 50% height | $PC_{50}$ | $\sum_{i=1}^N I(p_i(z) \geq \text{percentile}(p_i(z), 50))$ |
| Point count above 25% height | $PC_{25}$ | $\sum_{i=1}^N I(p_i(z) \geq \text{percentile}(p_i(z), 25))$ |

$Eucl(A, B)$ corresponds to Euclidean distance between points A and B.
$V_i^P$ is the number of points in the $i^{th}$ voxel cell, and $I$ is the indicator function

### 3.3 Plant Traits Characterization

Leaf nitrogen concentration and shoot biomass modeling were performed using three machine learning regression modeling techniques including Support Vector Regression (SVR), Random Forest Regression (RF) and Lasso Regression (Lasso), using the RGB spectral features and the LiDAR structural features defined in Table 1 and 2, respectively. The SVR model was used with the linear kernel [38] to accurately model the plant parameters, for its exception ability to model underlying relationship and quantifying feature relevance. Equation (2) shows the objective function and constraints of SVR with the linear kernel.

Let *x*_*k*_ *∈* ℝ^𝔽^ be *F* dimensional feature vector of the *k*^*th*^ training sample, *y*_*k*_ be the target value corresponding to the *k*^*th*^ training sample, *w* = [*w*_1_, *w*_2_, …, *w*_*F*_ ] be the vector containing the scalar weights associated with individual features in the feature set, *b* be the bias term, *ϵ* be the margin of tolerance, 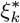 and *ξ*_*k*_ be the slack variables associated with each data point, and *C* be the regularization factor. The SVR is mathematically formulated as

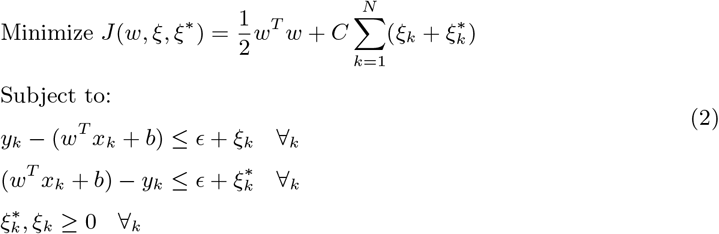

Here, the parameters *C* and *ϵ* were estimated using grid search. The value of *C* was varied over the range [0.01, 0.1, 1, 10, 100], while *ϵ* was varied over the range [0.01, 0.1, 0.5, 1], employing 5-fold cross-validation to assess model performance. The model fitting was repeated 100 times using different random sets of training and testing data, and the individual parameter estimates *ŷ*_*k*_ = (*w*^*T*^ *ϕ*(*x*_*k*_) + *b*) were averaged to obtain a reliable target parameter estimate.

The RF regression [39] follows an ensemble learning approach, where a collection of decision tree regression models operates independently to generate parameter estimates. The predictions from these individual trees are then averaged to produce a more reliable estimate. If, the RF model consists of *T* individual decision tree regression functions *f*_*t*_(*x*) where *t* = 1, 2, …, *T*, the target parameter estimate *ŷ*_*k*_ is calculated as the average of estimates obtained from the individual trees.

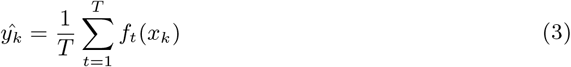

Here, the *T* and tree depth *D* of each tree were estimated using grid search on *T* for the range [1, 200] with a step size of 10, and *D* for the range [3, 5, 7, 10, 15] utilizing 5-fold cross-validation with, to assess the model performance.

We also model the plant parameters using the Lasso Regression [40] which is a widely used linear regression technique that incorporates feature regularization. Let, *y*_*k*_ is the true target variable value, *X*_*k*_ be the vector of predictors for the *k*^*th*^ observation, *w* be the vector of coefficients to be estimated, and *λ >* 0 be the regularization parameter that controls the strength of the penalty. Then, Lasso Regression is mathematically represented as

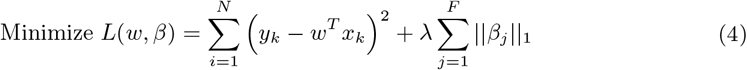

The *λ* was set by performing grid search on the range [0.01,0.1,1,10,100]. Once (4) is optimized, the target variable estimate for test data points *t* is obtained as *ŷ*_*t*_ = *w*^*T*^ *x*_*t*_.

For all the regression frameworks, the total available labeled data were divided in to training (70%) and testing (30%) sets. The accuracy assessment of the regression models was quantified using the Mean Error (ME), Mean Absolute Error (MAE), Root Mean Squared Error (RMSE), Relative Root Mean Squared Error (rRMSE), and Coefficient of Determination (R^2^) matrices, using the equations below.

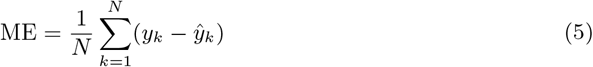

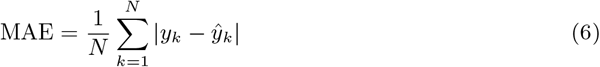

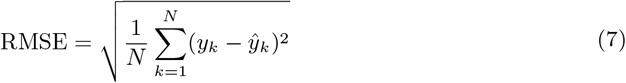

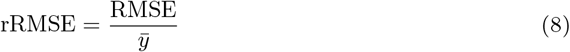

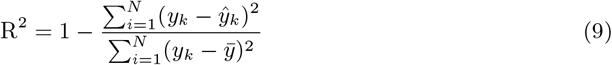

## 4 Results and Discussion

Fifteen RGB spectral features for individual Chinese spinach plants were derived from RGB leaf data, segmented using the MaskCNN algorithm, which achieved a high Intersection over Union (IoU) ratio of 0.92. Additionally, nine LiDAR structural features were estimated from the plant point cloud segment identified through a two-class K-means clustering algorithm. Variations in nitrogen dosing affected all 24 features, each serving as a proxy for specific plant traits or overall stress indicators. For example, RGB spectral features such as the Visible Atmospherically Resistant Index (VARI) and the Modified Green-Red Vegetation Index (MGRVI)-proxies for chlorophyll concentration and greenness—initially increased with nitrogen dosage but declined beyond a certain level. In contrast, indices like the Excess Red (ExR) and the Red-Green-Blue Vegetation Index (RGBVI), which are associated with plant stress, displayed an inverse relationship with nitrogen dosage. Figure 5 illustrates the effect of nitrogen dosage on the RGB spectral and the LiDAR structural feature values.

**Figure 5:**
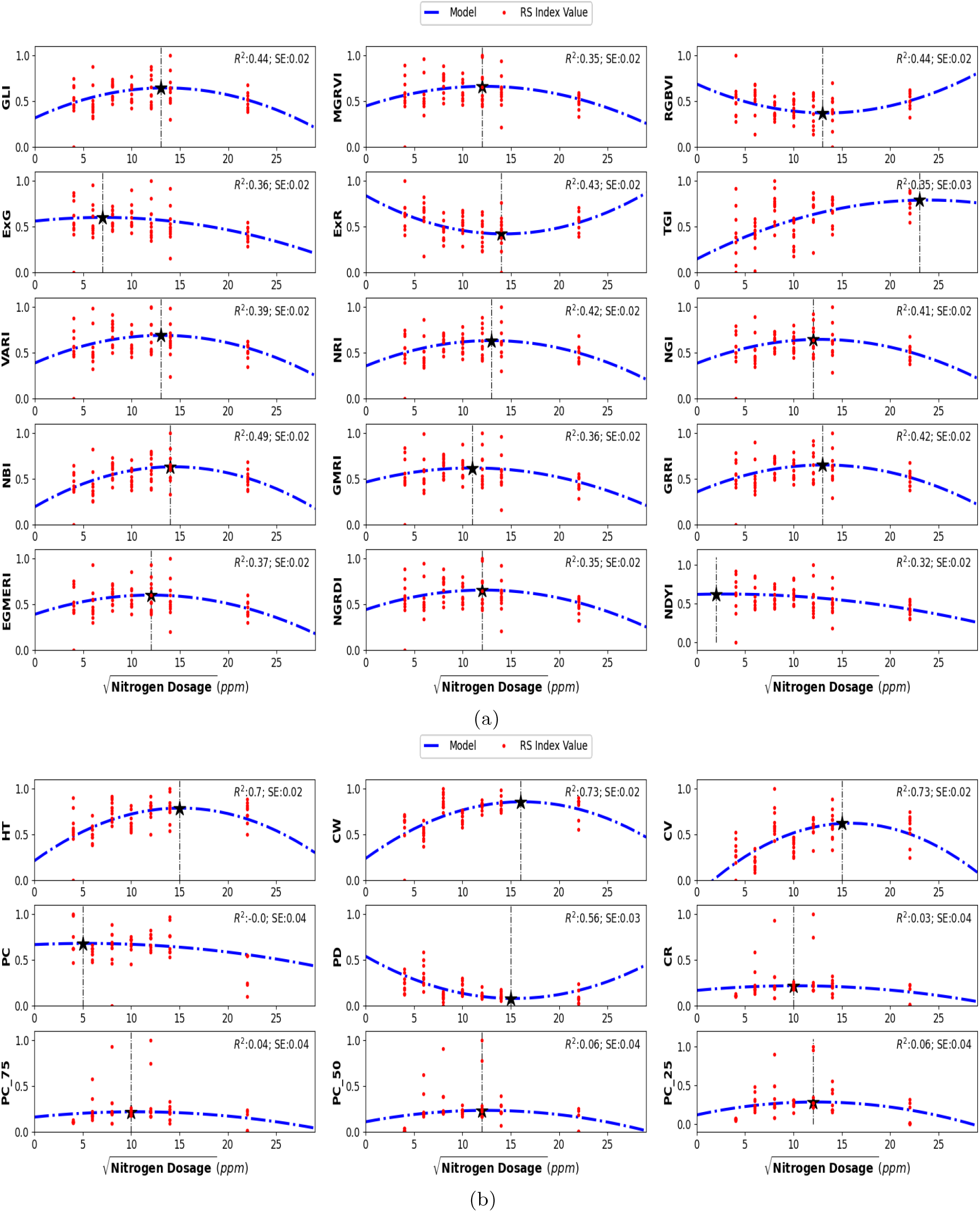
Second-order polynomial regression models illustrating the non-linear relationship between (a) RGB spectral features and (b) LiDAR structural features with nitrogen dosage in leaves. A square-root transformation was applied to the independent variable to normalize the initially skewed distribution. The minimum and maximum index values in each plot are indicated by the vertical dashed line.

The visual assessments in Figure 6 provide further insight into the effect of nitrogen treatments. Specifically, plants treated with both the lowest and highest nitrogen dosages exhibited reduced yields in terms of plant height and biomass, suggesting that extreme nitrogen levels may inhibit growth and development in Chinese spinach.

**Figure 6:**
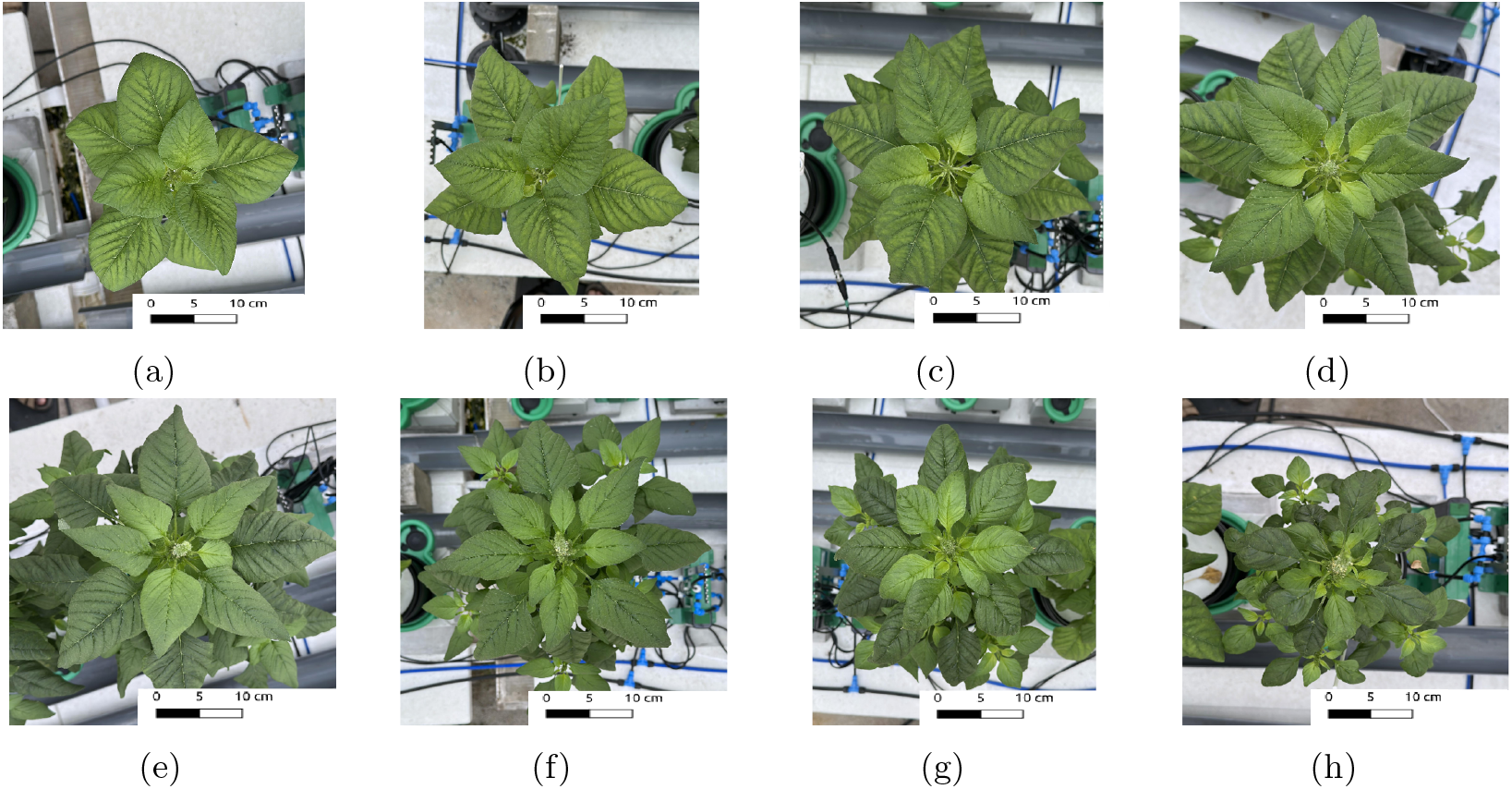
RGB images of Chinese spinach plants treated with varying nitrogen dosages: (a) 20 ppm, (b) 40 ppm, (c) 80 ppm, (d) 120 ppm, (e) 160 ppm, (f) 200 ppm, (g) 400 ppm, and (h) 800 ppm. Scale bars represent 10 cm.

The relationship between nitrogen dosing and individual RGB and LiDAR features was modeled using various functions, including linear, second-order, and third-order polynomials. The second-order polynomial model provided the best fit, effectively capturing the non-linear relationships between features and nitrogen dosage. To address the initially skewed distribution of nitrogen dosages, a square-root transformation was applied, which normalized the distribution and improved the linearity of the modeled relationships. This transformation enhanced the accuracy of plant parameter estimation based on smartphone-derived data. Figure 5 displays second-order polynomial models (dotted blue lines) that illustrate the relationships between various smartphone-derived indices and nitrogen dosages.

Combining RGB spectral and LiDAR structural features offered complementary information on leaf nitrogen concentration and shoot biomass, thereby improving the precision and reliability of crop trait monitoring—especially in indoor greenhouse environments for crops like Chinese spinach. This complementarity is further evidenced in Figure 7, where low correlations are observed between the RGB spectral and LiDAR structural feature sets, while higher correlations are found within each feature set. This finding indicates that RGB imagery and LiDAR data provide unique, complementary insights.

**Figure 7:**
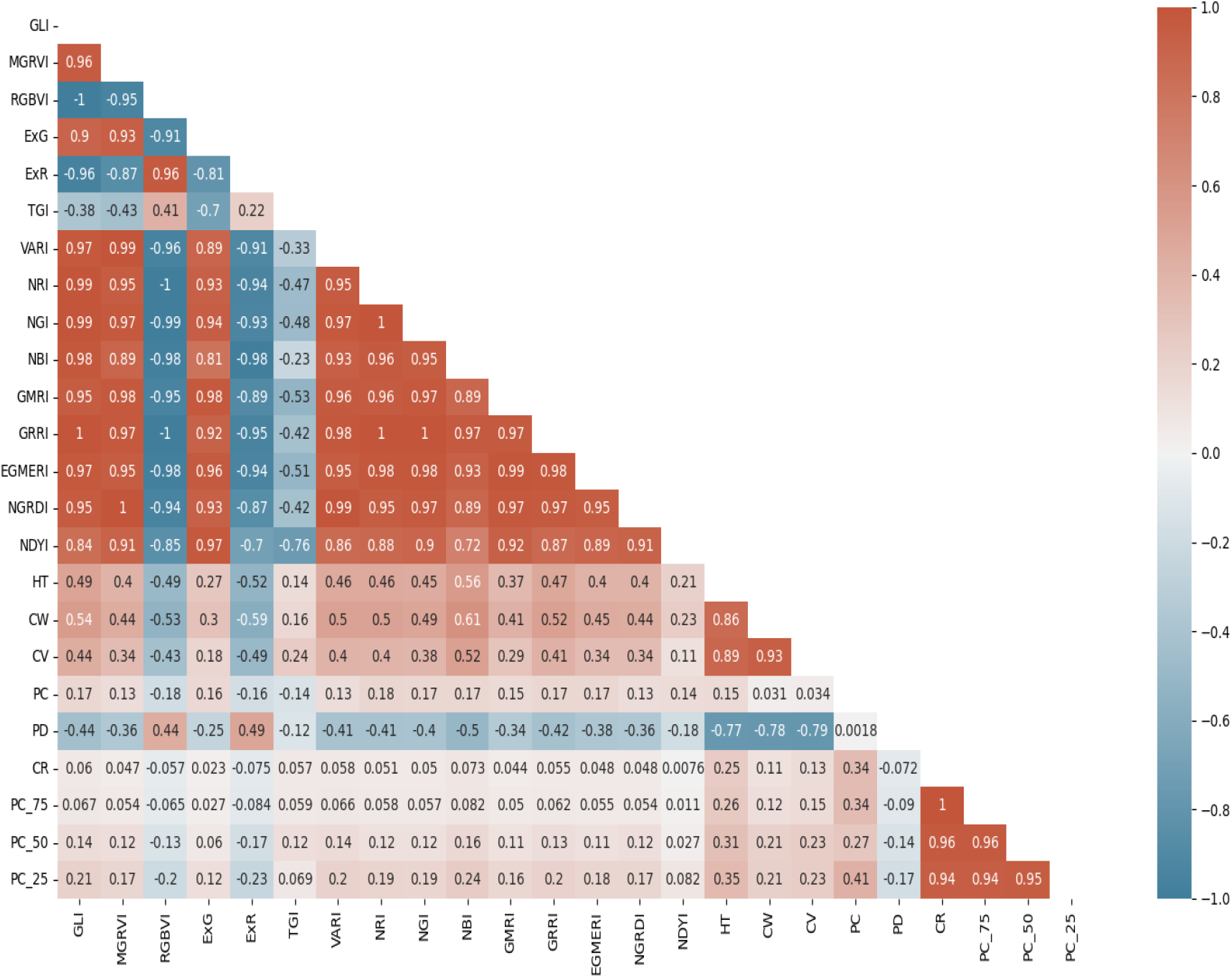
Correlation matrix of RGB spectral indices and LiDAR structural indices derived from smartphone RGB imagery and LiDAR data. The matrix illustrates the relationships between different indices, with color intensity indicating the strength and direction of the correlation (red for positive, blue for negative correlations).

To model three key plant parameters—leaf *TRN* concentration, leaf *NO*_3_ concentration, and shoot *DW* biomass-we employed three machine learning algorithms: SVR, RF, and Lasso Regression. Modeling was conducted using three distinct feature-set configurations: RGB spectral features only, LiDAR structural features only, and A combination of both RGB spectral and LiDAR structural features. In the case *TRN* modeling, the optimal *C* and *ϵ* for SVR were estimated as 10 and 0.1, respectively, *T* and *D* parameters of RF were estimated to be 100 and 5 respectively, and *λ* was estimated to be 0.1 in the case of lasso regression. This multi-configuration approach allowed us to evaluate the unique contributions of each feature set and their combinations in predicting key plant parameters. Across all feature-set configurations, the RF model consistently performed well, yielding relatively lower Root Mean Squared Error (RMSE) and higher *R*^2^ values compared to SVR and Lasso, particularly when using the combined feature set. Lasso Regression, however, showed high Mean Error (ME) and Mean Absolute Error (MAE) for the structural feature set (e.g., ME of 12.53 g), indicating possible overfitting or instability in Lasso’s performance for DW estimation.

Additionally, a clear pattern emerged across the different feature sets: for *TRN* and *NO*_3_ estimation, the RGB feature set yielded the lowest prediction errors, while for shoot *DW*, the structural feature set provided the most accurate estimates with the lowest errors. Overall, the results demonstrate the effectiveness of combining RGB imagery and LiDAR data in crop parameter modeling and highlight the robustness of the RF algorithm in managing complex multivariate data. The model estimation performance for *TRN* concentration, *NO*_3_ concentration, and shoot *DW* - quantified using ME, MAE, RMSE, rRMSE, and *R*^2^ values—is provided in Table 3 and Table 4.

**Table 3:**
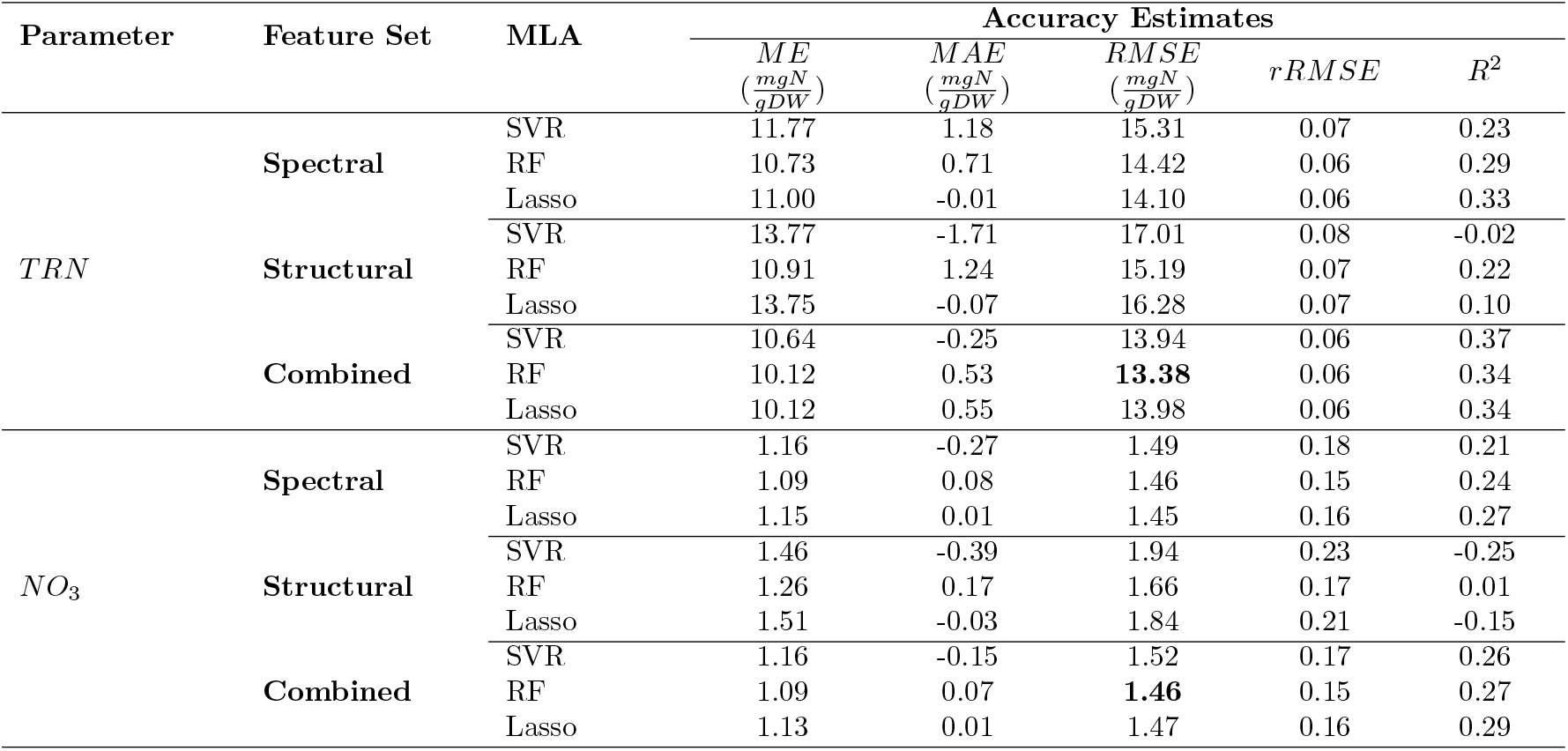
The *ME, MAE, RMSE, rRMSE*, and *R*^2^ for leaf nitrogen concentration estimations done using the SVR, the RF, and the Lasso Regression.

| Parameter | Feature Set | MLA | Accuracy Estimates |  |  |  |  |
| --- | --- | --- | --- | --- | --- | --- | --- |
| | | | *ME*<br>($\frac{mgN}{gDW}$) | *MAE*<br>($\frac{mgN}{gDW}$) | *RMSE*<br>($\frac{mgN}{gDW}$) | *rRMSE* | *R<sup>2</sup>* |
| *TRN* | Spectral | SVR | 11.77 | 1.18 | 15.31 | 0.07 | 0.23 |
|  |  | RF | 10.73 | 0.71 | 14.42 | 0.06 | 0.29 |
|  |  | Lasso | 11.00 | -0.01 | 14.10 | 0.06 | 0.33 |
|  | Structural | SVR | 13.77 | -1.71 | 17.01 | 0.08 | -0.02 |
|  |  | RF | 10.91 | 1.24 | 15.19 | 0.07 | 0.22 |
|  |  | Lasso | 13.75 | -0.07 | 16.28 | 0.07 | 0.10 |
|  | Combined | SVR | 10.64 | -0.25 | 13.94 | 0.06 | 0.37 |
|  |  | RF | 10.12 | 0.53 | **13.38** | 0.06 | 0.34 |
|  |  | Lasso | 10.12 | 0.55 | 13.98 | 0.06 | 0.34 |
| *NO<sub>3</sub>* | Spectral | SVR | 1.16 | -0.27 | 1.49 | 0.18 | 0.21 |
|  |  | RF | 1.09 | 0.08 | 1.46 | 0.15 | 0.24 |
|  |  | Lasso | 1.15 | 0.01 | 1.45 | 0.16 | 0.27 |
|  | Structural | SVR | 1.46 | -0.39 | 1.94 | 0.23 | -0.25 |
|  |  | RF | 1.26 | 0.17 | 1.66 | 0.17 | 0.01 |
|  |  | Lasso | 1.51 | -0.03 | 1.84 | 0.21 | -0.15 |
|  | Combined | SVR | 1.16 | -0.15 | 1.52 | 0.17 | 0.26 |
|  |  | RF | 1.09 | 0.07 | **1.46** | 0.15 | 0.27 |
|  |  | Lasso | 1.13 | 0.01 | 1.47 | 0.16 | 0.29 |

**Table 4:**
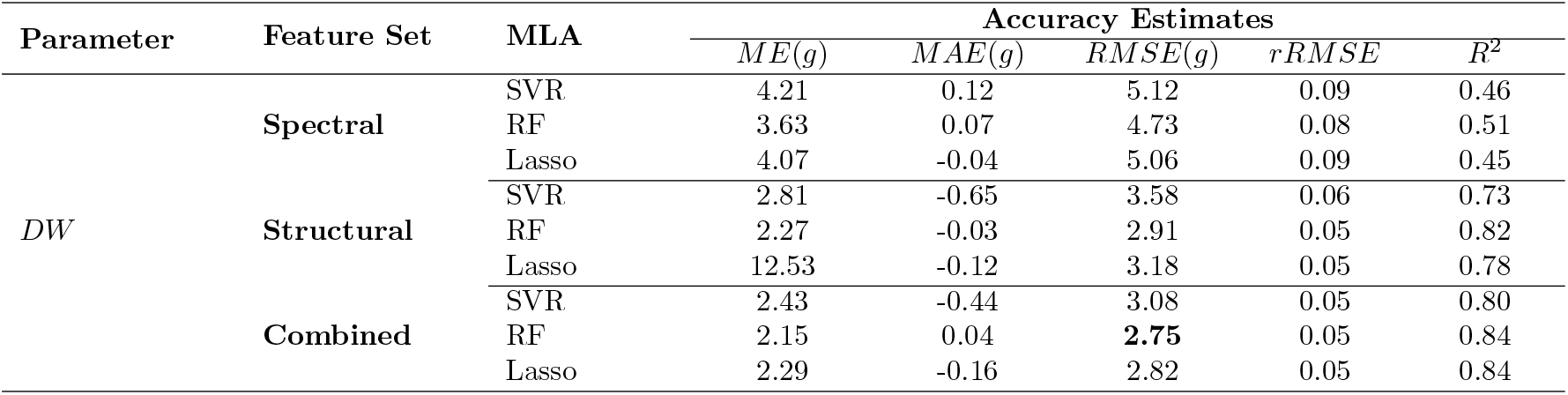
The *ME, MAE, RMSE, rRMSE*, and *R*^2^ for shoot *DW* estimation done using the SVR, the RF, and the Lasso Regression.

| Parameter | Feature Set | MLA | Accuracy Estimates |  |  |  |  |
| --- | --- | --- | --- | --- | --- | --- | --- |
|  |  |  | *ME(g)* | *MAE(g)* | *RMSE(g)* | *rRMSE* | *R<sup>2</sup>* |
| *DW* | Spectral | SVR | 4.21 | 0.12 | 5.12 | 0.09 | 0.46 |
|  |  | RF | 3.63 | 0.07 | 4.73 | 0.08 | 0.51 |
|  |  | Lasso | 4.07 | -0.04 | 5.06 | 0.09 | 0.45 |
|  | Structural | SVR | 2.81 | -0.65 | 3.58 | 0.06 | 0.73 |
|  |  | RF | 2.27 | -0.03 | 2.91 | 0.05 | 0.82 |
|  |  | Lasso | 12.53 | -0.12 | 3.18 | 0.05 | 0.78 |
|  | Combined | SVR | 2.43 | -0.44 | 3.08 | 0.05 | 0.80 |
|  |  | RF | 2.15 | 0.04 | **2.75** | 0.05 | 0.84 |
|  |  | Lasso | 2.29 | -0.16 | 2.82 | 0.05 | 0.84 |

Beyond plant parameter estimation, we also focused on determining the optimal nitrogen dosage to maximize plant biomass and vigor using smartphone-derived RGB and LiDAR data. The optimal nitrogen dosage is defined as the dosage level that maximizes or minimizes individual RGB spectral and LiDAR structural features, each serving as a proxy for specific plant traits, such as chlorophyll concentration, leaf senescence, and shoot biomass. Since all smartphone features were modeled using second-order polynomials, peak values of individual features were determined by calculating the second derivative of each polynomial and setting it to zero. This process allowed us to pinpoint the critical nitrogen dosage that optimizes each plant trait. In this study, the optimal nitrogen dosage for Chinese spinach was estimated to be 149.9 ppm. Notably, this value is close to the optimal dosage of 120.0 ppm derived from in-situ transpiration data collected using the PlantArray system, as reported in [5]. The agreement between the smartphone-derived optimal dosage and the dosage estimated through sophisticated transpiration data further validates the accuracy and reliability of our approach using smartphone RGB and LiDAR data. However, it is important to note that [5] did not investigate nitrogen dosages between 120.0 ppm and 160.0 ppm, making precise validation of the optimal nitrogen dosage challenging with the available data.

A detailed feature relevance analysis was conducted to assess the importance of various RGB and LiDAR-derived features in predicting key target variables: leaf *TRN* concentration, *NO*_3_ concentration, and *DW* biomass. Feature relevance was inferred based on the weights assigned to each feature during SVR model training. These weights were averaged across 100 modeling iterations, with each iteration using randomly generated training and test data sets to ensure robustness. The results indicate that feature importance varied significantly across models, reflecting the specific demands of each prediction task. For example, models developed to predict *TRN* and *NO*_3_ concentration assigned higher importance to the RGB features, likely because these spectral indices are closely related to nitrogen content, which affects reflectance in the visible spectrum. In contrast, models estimating shoot *DW* generally assigned greater weight to structural features obtained from LiDAR data, reflecting that biomass estimation can be more reliably modeled using plant morphology rather than spectral characteristics. Specifically, *DW* estimation showed high reliance on LiDAR-derived structural indices such as crown volume (*CV*) and point count at 75% height (*PC*_75_), which describe the plant’s physical structure and distribution. Conversely, for *TRN* and *NO*_3_ estimation, the models placed more emphasis on spectral features such as the Triangular Greenness Index (*TGI*) and Excess Green (*ExG*), both closely related to chlorophyll content and leaf greenness. These findings suggest a clear pattern: LiDAR-derived structural features are more effective for estimating morphological traits like biomass, while RGB features are better suited for predicting physiological traits like nitrogen and chlorophyll concentration. This distinction underscores the complementary nature of smartphone RGB imagery and LiDAR data in crop parameter modeling, with each sensor type contributing uniquely to different aspects of plant structure and function (Fig. 8).

**Figure 8:**
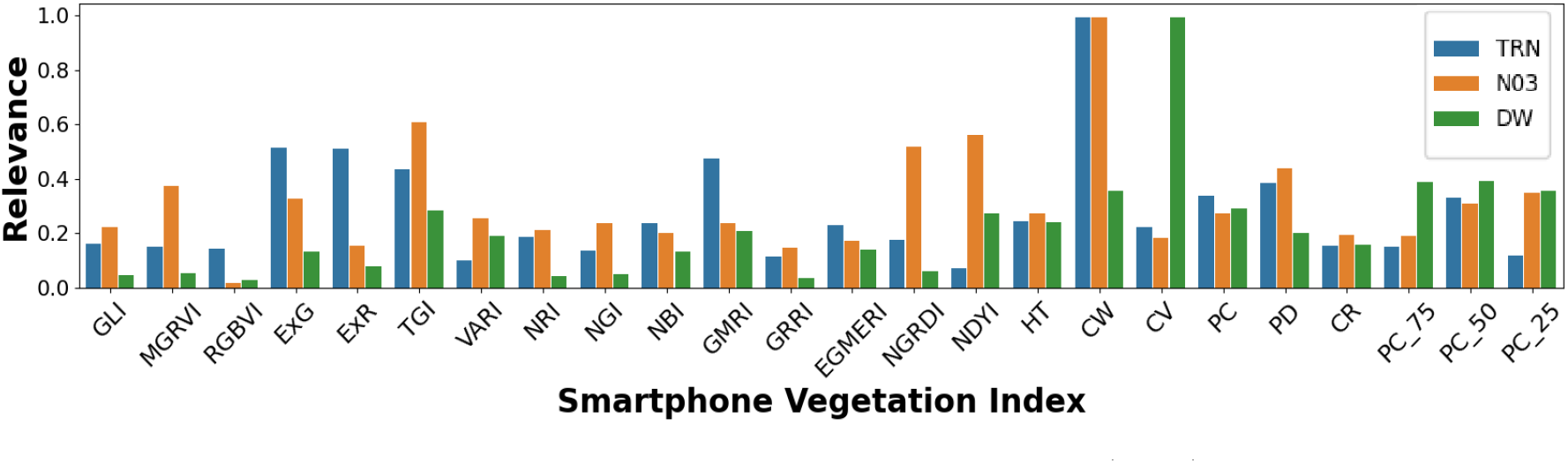
Normalized weights from the Support Vector Regression (SVR) model, indicating the relative importance of each RGB spectral and LiDAR structural feature in estimating leaf total reduced nitrogen (*TRN*), leaf nitrate (*NO*_3_), and shoot dry weight (*DW*).

To evaluate our methodology, we compared its performance with a leading state-of-the-art leaf nitrogen concentration estimation approach (SoA-NC) that relies solely on smartphone RGB imagery. This benchmark method uses the Dark Green Color Index (*DGCI*), derived from intensity, hue, and saturation metrics of the plant leaves [41]. However, using only a single RGB image feature index limits its accuracy in estimating leaf nitrogen concentration. Our results illustrate the superior performance of our models, which consistently produced more accurate and precise estimates of leaf *TRN* and *NO*_3_ concentrations compared to the SoA-NC method. This improvement highlights the advantage of integrating advanced machine learning algorithms with both spectral and structural features derived from smartphone RGB and LiDAR sensors. Unlike the SoA-NC method, which focuses solely on spectral data, our approach leverages the complementary strengths of RGB features (e.g., color indices) and LiDAR structural features (e.g., plant height and canopy shape). Whatsoever, high variance in individual feature index values, even within a nitrogen treatment group, due to non-uniform lighting or shading on plant leaves, is a known source of error, which manifests as significant variance in *TRN, NO*_3_ and *DW* estimates.

We also compared shoot *DW* estimation performance with a state-of-the-art leaf-area-based shoot *DW* estimation (SoA-DW) method [42]. The SoA-DW method estimates shoot *DW* by: (a) segmenting plant leaves using Otsu’s thresholding on the green channel, (b) calculating leaf area (*LA*_*p*_) as the product of the total number of plant pixels (*PPN*) in the image, the individual pixel length (*PL*) of the phone camera, and the magnification factor (*MF*) of the phone camera, and (c) estimating shoot *DW* using a linear model based on *LA*_*p*_ and actual *DW* values obtained from destructive sampling. Our proposed shoot *DW* estimation method more accurately tracked actual shoot *DW* compared to the SoA-DW method. The reduced performance of the SoA-*DW* method likely results from its inability to capture plant height from images and from errors in leaf delineation due to non-uniform lighting and shading.

A direct comparison between the *TRN, NO*_3_, and *DW* estimates obtained from our non-destructive model and those derived from traditional destructive sampling methods is also provided to further validate the accuracy and reliability of our methodology (Fig. 9). The results demonstrate that the proposed handheld smartphone-based system can accurately estimate leaf *TRN*, leaf *NO*_3_, and shoot *DW* without the need for manual harvesting, outperforming state-of-the-art methods that rely solely on RGB data.

**Figure 9:**
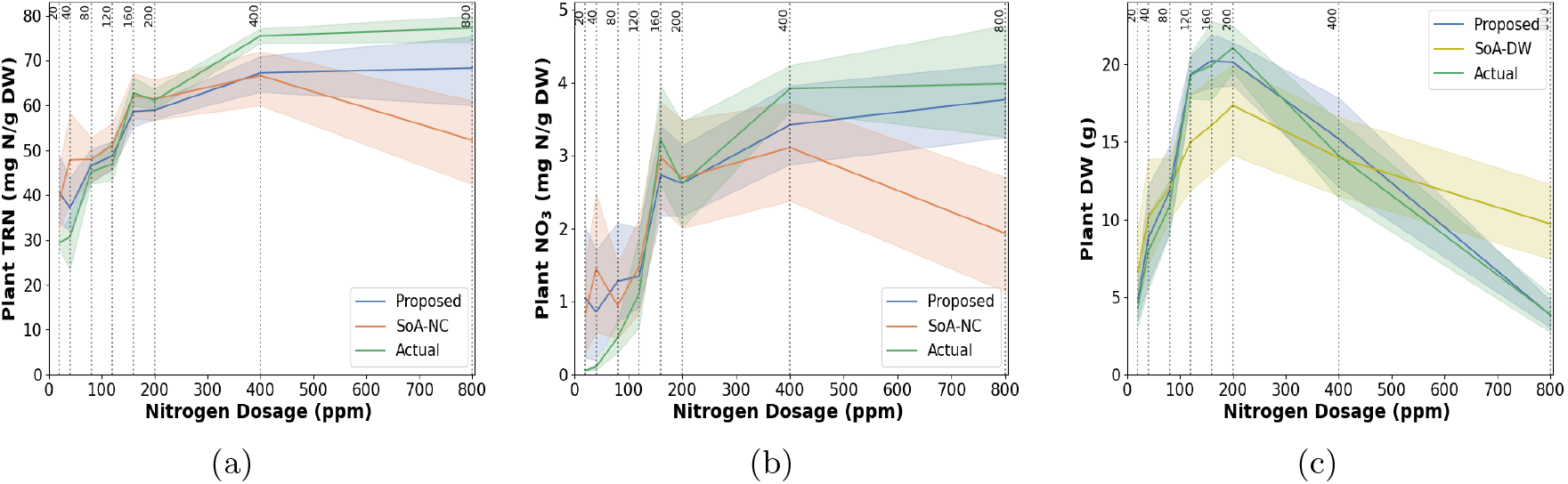
Model estimation performance for (a) plant total reduced nitrogen (*TRN*), (b) nitrate (*NO*_3_), and (c) dry weight (*DW*) in Chinese spinach across different nitrogen dosages. Results are shown for the proposed method, state-of-the-art (SoA) nitrogen concentration (SoA-NC) and dry weight (SoA-DW) estimation approaches, with reference data obtained through destructive leaf analysis.

## 5 Summary and Conclusion

Smartphone-based RGB imagery and LiDAR data show strong potential for modeling the biophysical and structural parameters of *Amaranthus dubius* (Chinese spinach), serving as a case study for subtropical leafy vegetable crops. The combined use of spectral and structural features derived from smartphone RGB and LiDAR data, integrated with Random Forest regression, enables accurate modeling of leaf nitrogen concentration and shoot dry-weight biomass in Chinese spinach. Experimental results indicate that smartphone-derived RGB and LiDAR data can reliably estimate leaf total reduced nitrogen (*TRN*), nitrate (*NO*_3_), and dry weight (*DW*) with average RMSE values as low as 3.9 mg N/g DW, 1.46 mg N/g DW, and 2.7 mg, respectively. Spectral indices were found to be more effective for estimating leaf nitrogen concentration, whereas structural indices provided better performance for shoot biomass estimation. The optimal nitrogen dosage for maximizing yield and fertilizer use efficiency in Chinese spinach was determined to be 149.9 ppm, based on RGB spectral and LiDAR structural features. Future research will focus on expanding the application of this methodology to assess plant requirements for other essential mineral nutrients. The promising results from this study suggest strong potential for industrial-scale implementation, leveraging affordable and accessible smartphone technology. Integration of the approach into automated robotic systems such as gantries and drones has the potential to enable real-time data capture and analysis, providing farmers with convenient, precise insights to optimize fertilizer usage and crop health.

## Acknowledgement

The Israel-Singapore Urban Farm (iSURF) is supported by the National Research Foundation, Prime Minister’s Office, Singapore, under its Campus for Research Excellence And Technological Enterprise (CREATE) programme [NRF2020-THE003-0008]. This work was supported in part by the Hebrew University of Jerusalem Center for Interdisciplinary Data Science Research (CIDR). We gratefully acknowledge assistance from iSURF program collaborators Netatech Pte Ltd, Singapore and Mr. David Tan.

